# KipOTIA detoxifies 5-oxoproline and promotes the growth of *Clostridioides difficile*

**DOI:** 10.1101/2024.05.01.592088

**Authors:** Cheyenne D. Lee, Arshad Rizvi, Shonna M. McBride

**Affiliations:** Department of Microbiology and Immunology, Emory University School of Medicine, Emory Antibiotic Resistance Center, Atlanta, GA, USA

**Keywords:** *Clostridium difficile*, kinase inhibitory protein, 5-oxoproline, defined minimal medium, Niehaus defined medium, pxp

## Abstract

*Clostridioides difficile* is an anaerobic enteric pathogen that disseminates in the environment as a dormant spore. For *C. difficile* and other sporulating bacteria, the initiation of sporulation is a regulated process that prevents spore formation under favorable growth conditions. In *Bacillus subtilis*, one such mechanism for preventing sporulation is the Kinase Inhibitory Protein, KipI, which impedes activation of the main sporulation kinase. In addition, KipI functions as part of a complex that detoxifies the intermediate metabolite, 5-oxoproline (OP), a harmful by-product of glutamic acid. In this study, we investigate the orthologous Kip proteins in *C. difficile* to determine their roles in the regulation of sporulation and metabolism. Using deletion mutants in *kipIA* and the full *kipOTIA* operon, we show that unlike in *B. subtilis,* the Kip proteins have no significant impact on sporulation. However, we found that the *kip* operon encodes a functional oxoprolinase that facilitates detoxification of OP. Further, our data demonstrate that KipOTIA not only detoxifies OP, but also allows OP to be used as a nutrient source that supports the robust growth of *C. difficile*, thereby facilitating the conversion of a toxic byproduct of metabolism into an effective energy source.

## INTRODUCTION

*Clostridioides difficile* is an anaerobic Gram-positive pathogen that causes illnesses ranging from abdominal pain and diarrhea, to life-threatening pseudomembranous colitis (Dubberke and Olsen, 2012; Finn et al., 2021). *C. difficile* infections are easily spread and difficult to contain because the spore form of the bacterium is resistant to disinfectants and allows the pathogen to survive nearly indefinitely in a dormant state (Ali et al., 2011; A.N. Edwards et al., 2016; Shen, 2020). Although the structure and general form of *C. difficile* endospores are similar to other species, the environmental and nutritional conditions that lead to sporulation vary greatly among spore-forming bacteria (Lee et al., 2022; Mehdizadeh Gohari et al., 2024; Shen et al., 2019). While some basic nutritional requirements and regulatory factors are shared among the sporulating Firmicutes, it is not possible to predict the role of a given factor based on the presence of orthologs with a known function in another spore-former (A. N. Edwards et al., 2016; Edwards et al., 2022, 2014).

Much of what is known about sporulation and nutrition in Gram-positive bacteria is based on extensive studies performed in the model spore-forming species, *Bacillus subtilis* (Errington, 2003; Sonenshein, 2007, 2000). In *B. subtilis*, sporulation initiates with the accumulation of the activated (phosphorylated) master regulator, Spo0A. The activation of Spo0A in *B. subtilis* is subject to a succession of negative and positive regulatory inputs that control the flow of phosphoryl groups to Spo0A via a phosphorelay. A key driver of the sporulation phosphorelay is the sensor kinase, KinA, which can be inhibited by the anti-kinase KipI, resulting in decreased sporulation (Wang et al., 1997). In turn, the co-transcribed factor, KipA, binds to KipI to prevent KipI interference with KinA activation of the phosphorelay (Jacques et al., 2008, 2011). Though the functions of KipI and KipA are known for *B. subtilis*, their roles in the sporulation regulation of other species have not been described.

While sporulation regulation is one function of the KipI and KipA proteins in *B. subtilis*, other studies have demonstrated metabolic functions of these proteins (Niehaus et al., 2017). In *B. subtilis*, *Escherichia coli*, and *Pseudomonas putida*, KipI and KipA bind with KipO (also known as PxpA,B,C) to form an enzyme that metabolizes 5-oxoproline (Li et al., 1988; Niehaus et al., 2017; Seddon et al., 1984; Seddon and Meister, 1986). 5-oxoproline (OP), or L-pyroglutamic acid, is a ubiquitous waste product formed in bacteria from the spontaneous cyclization of glutamate, glutamine, and ɣ-glutamyl phosphate (Niehaus et al., 2017; Seddon et al., 1984). Free 5-oxoproline is also formed from the action of the pyrrolidone carboxyl peptidase (Pcp) which cleaves N-terminal 5-oxoproline from cyclized glutaminyl or glutamyl on peptides and is found in many prokaryotic organisms (Awadé et al., 1992, 1994; Doolittle and Armentrout, 1968). Accumulation of 5-oxoproline is toxic to eukaryotic and prokaryotic cells and impairs growth, but organisms that encode KipOIA orthologs can convert OP to glutamic acid, thereby preventing OP toxicity (Kumar and Bachhawat, 2010; Park et al., 2001; Vetter et al., 2019).

In this study, we sought to determine whether the *C. difficile* Kip orthologs function as regulators of sporulation and/or metabolism of 5-oxoproline. Our results demonstrate that, unlike *B. subtilis*, KipI and KipA in *C. difficile* do not play a major role in sporulation initiation. We also demonstrate that the *kipOTIA* operon encodes a functional oxoprolinase that allows for growth in OP. Further, our results indicate that 5-oxoproline is not only detoxified by *C. difficile*, it is an effective and valuable nutrient source. Overall, our data indicate that the Kip proteins function primarily as a 5-oxoprolinase that promotes the growth of *C. difficile* in the presence of this toxic metabolite.

## RESULTS

### Identification and disruption of the KipOTIA orthologs in *C. difficile*

The putative KipI and KipA proteins of *C. difficile* were identified in the strain 630 genome by sequence similarity to the Kip proteins of *B. subtilis* (**Figure S1A**). The *C. difficile* KipI and KipA are encoded immediately downstream of orthologs of *kipO* and *kipT* (CD630_13840-13870), which are also found within the *B. subtilis kip* locus (**Figure S1A**). *C. difficile* does not encode three additional genes (*pxpI, kipR*, and *lipC*) that are encoded within the *B. subtilis kip* region. We examined transcription of the *C. difficile kip* genes in the region to determine if they were encoded as an operon. We performed nested PCR on *C. difficile* cDNA templates and demonstrated that *kipOTIA* are transcribed as a polycistronic unit (**Figure S1B**). Based on the similarities between the shared proteins, we hypothesized that the *C. difficile* Kip proteins may function as regulators of sporulation and metabolism, as observed in *B. subtilis*.

### KipOTIA does not inhibit sporulation in *C. difficile*

To assess the Kip protein functions in *C. difficile*, we generated Δ*kipIA* (MC1903) and Δ*kipOTIA* (MC2375) mutants by allelic exchange and replacement with an *ermB* cassette. As controls, both mutants were complemented with the *kipOTIA* operon at the *kip* locus (**Table S1, Figure S2**) and all resultant strains were verified by whole-genome sequencing. As KipI is an inhibitor of sporulation in *B. subtilis*, we next examined the impact of deleting the *kip* genes on *C. difficile* sporulation. Sporulation assays were performed on 70:30 sporulation agar, comparing the Δ*kipIA* and Δ*kipOTIA* mutants to the parent strain (**Figure 1**). Deletion of *C. difficile kipIA* resulted in a modest decrease in sporulation, while deletion of the entire *kipOTIA* operon had no apparent effects. The decreased sporulation for Δ*kipIA* mutant, but not the Δ*kipOTIA* mutant, suggests that expression of *kipO* and *kipT* without *kipI* and *kipA*, is detrimental to spore formation. The *kip* deletion phenotypes in *C. difficile* are in contrast to *B. subtilis*, where *kipIA* mutants produce more spores and over-expression of *kipI* reduces sporulation (Wang et al., 1997). These results strongly suggest that unlike in *B. subtilis,* KipI does not inhibit a sporulation-specific kinase in *C. difficile*.

**Figure 1.**
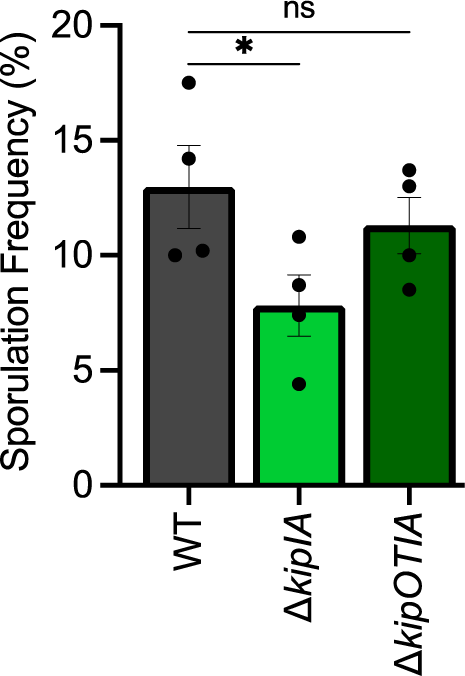
KipOTIA does not inhibit *C. difficile* sporulation. Ethanol-resistant sporulation frequencies of *C. difficile* strain 630Δ*erm* (WT), Δ*kipIA* mutant (MC1903), and the Δ*kipOTIA* mutant (MC2375) grown for 24 h on 70:30 sporulation agar. The means and SEM of four independent experiments are shown. Data were analyzed by one-way ANOVA with Dunnett’s multiple comparison test; \**P* <0.05

To better understand the underlying basis for the Δ*kipIA* sporulation decrease in *C. difficile*, we examined gene expression in sporulation medium for the Δ*kipIA* mutant relative to the parent strain by RNA-seq (**Table 1**; MC1970, Δ*kipIA* pMC123 and MC324, 630Δ*erm* pMC123). Only 25 genes were differentially regulated more than 2-fold in the Δ*kipIA* mutant, but the greatest transcriptional differences were observed for the three genes representing the oxidative branch of the *C. difficile* TCA cycle (*acnB, aksA,* and *icd*), suggesting changes in metabolism or redox within the mutant (Neumann-Schaal et al., 2019). Of the remaining transcriptional changes, more than half encode proteins of unknown function and two were sporulation-related genes. Aside from the changes in TCA gene expression, no patterns signifying specific regulatory factors, pathways, or processes were evident.

**Table 1.** Genes differentially expressed in a Δ*kipIA* mutant.

| Gene<sup>a</sup> | Fold Change<br>$\Delta kiplA/WT^b$ | Product | P value |
| --- | --- | --- | --- |
| **Increased in $\Delta kiplA$** |  |  |  |
| CD630_08330 | 4.66 | AcnB | 5.7E-17 |
| CD630_08320 | 4.57 | Re-citrate synthase | 2.4E-15 |
| CD630_08340 | 4.16 | lcd | 2.6E-13 |
| CD630_06170 | 2.38 | membrane protein | 1.6E-07 |
| CD630_04120 | 2.34 | conjugative transfer protein | 1.1E-05 |
| CD630_06160 | 2.18 | MerR family regulator | 5.4E-05 |
| CD630_05300 | 2.09 | membrane protein | 1.1E-06 |
| CD630_36160 | 2.08 | potassium/proton antiporter | 7.2E-06 |
| CD630_17170 | 2.05 | hypothetical protein | 1.5E-07 |
| CD630_02030 | 2.02 | UvrABC system protein A 1 | 1.8E-06 |
| CD630_07410 | 2.02 | GlpK1 | 0.0017 |
| **Decreased in $\Delta kiplA$** |  |  |  |
| CD630_13010 | 0.36 | membrane protein | 4.1E-06 |
| CD630_14182 | 0.41 | hypothetical protein | 7.2E-08 |
| CD630_05720 | 0.42 | sporulation protein | 5.1E-10 |
| CD630_07121 | 0.43 | hypothetical protein | 1.2E-04 |
| CD630_11990 | 0.43 | SpolIIAH | 3.3E-04 |
| CD630_34570 | 0.44 | hypothetical protein | 3.8E-04 |
| CD630_17281 | 0.44 | hypothetical protein | 2.4E-04 |
| CD630_17900 | 0.46 | exonuclease | 5.3E-05 |
| CD630_24570 | 0.47 | hypothetical protein | 2.3E-04 |
| CD630_20250 | 0.49 | membrane protein | 3.6E-04 |
| CD630_26030 | 0.49 | CdtR | 2.8E-05 |
| CD630_09410 | 0.50 | hypothetical protein | 2.2E-05 |
| CD630_12371 | 0.50 | hypothetical protein | 0.0028 |
| CD630_19670 | 0.50 | hypothetical protein | 5.4E-04 |

<sup>a</sup>Gene accession numbers for strain 630 (GenBank accession no. NC\_009089.1). Listed genes represent those transcripts with a $\geq 2$ -fold change in expression and a *P* value of $\leq 0.05$ as determined by DESeq2 analysis, as described in Methods. Genes representing *kipl*, *kipA*, and the *erm* selectable marker are excluded.
<sup>b</sup>Ratio of expression in $\Delta kiplA$ pMC123 (MC1970)/630 $\Delta erm$ pMC123 (MC324), as determined by RNA sequencing analysis as described in the Methods.

Since the Kip complex did not inhibit sporulation in our standard assay, we considered that the Kip proteins may conditionally influence sporulation, as observed in *B. subtilis*. In *B. subtilis,* the *kip* operon is induced by glucose and repressed by glutamine (Wang et al., 1997). Previous studies in *C. difficile* found that the *kipOTIA* operon is repressed by proline through PrdR, and by glucose, partially through CcpA, but the impact of these factors on *kip* expression was modest (2-3 fold) (Antunes et al., 2012; Bouillaut et al., 2019). Data from pilot experiments suggested that the secondary bile salt deoxycholate (DCA) induced transcription of the *kipOTIA* genes. We further examined the effects of bile salts on expression of the *kip* operon by assessing *kipI* transcription. To this end, wild-type *C. difficile* was grown in BHIS broth supplemented with physiologically relevant concentrations of deoxycholate (DCA, 0.5 mM), chenodeoxycholate (CDCA, 0.5 mM), lithocholate (LCA, 0.05 mM), cholate (CA, 2 mM), taurocholate (TA, 10 mM), glycocholate (GCA, 10 mM), taurochenodeoxycholate (TCDCA, 10 mM), or glycochenodeoxycholate (GCDCA, 1.5 mM) (Northfield and McColl, 1973) and samples collected for qRT-PCR analysis. The hydrophobic bile salts DCA, CDCA, LCA, and GCDCA significantly increased *kipI* expression, while CA and TA did not (**Figure 2**). To determine if the detergent properties of bile salts played a role in *kipI* expression*, C. difficile* was also exposed to sub-inhibitory concentrations of the detergents Triton X-100 and sodium dodecyl sulfate (SDS) (**Figure 2**). Both detergents induced *kipI* expression, but the differences were not statistically significant. These data demonstrate that *kip* expression is robustly induced by several bile salts that impact growth, and that *kip* induction is not universal to all bile acids or detergents.

**Figure 2.**
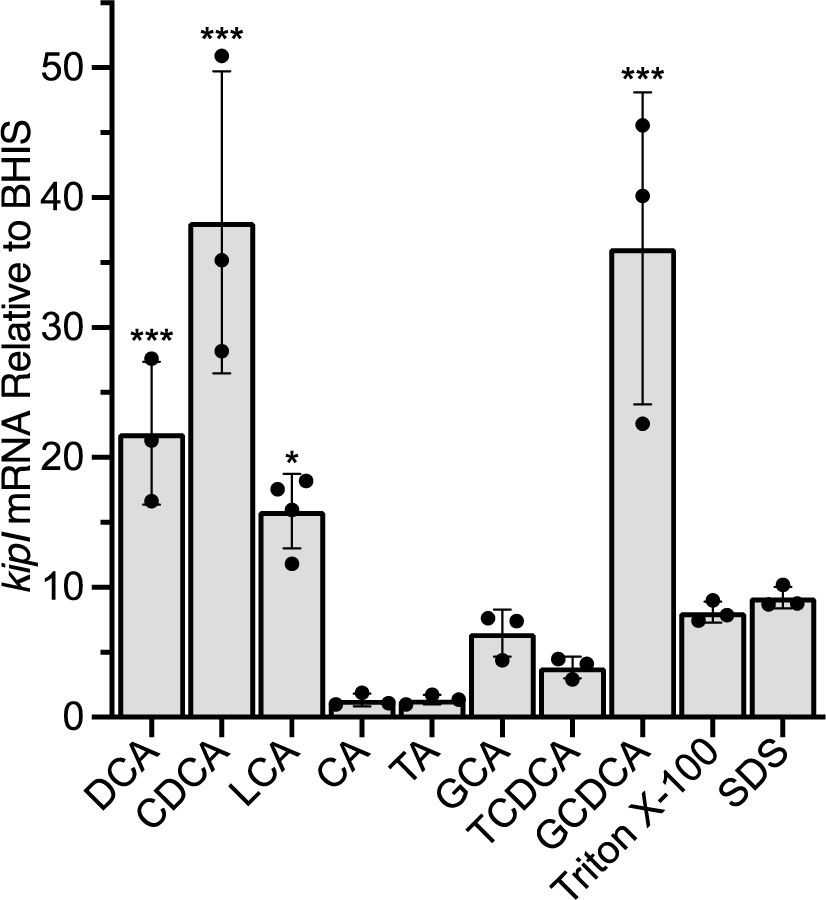
*C. difficile kipI* expression is induced by specific bile salts. WT *C. difficile* (630Δ*erm*) was grown in BHIS broth +/- physiological concentrations of bile salts, or detergents, as follows: deoxycholate (DCA, 0.5 mM), chenodeoxycholate (CDCA, 0.5 mM), lithocholate (LCA, 0.05 mM), cholate (CA, 2 mM), taurocholate (TA, 10 mM), glycocholate (GCA, 10 mM), taurochenodeoxycholate (TCDCA, 10 mM), glycochenodeoxycholate (GCDCA, 1.5 mM), 0.008% triton X-100, or 0.0045% sodium dodecyl sulfate (SDS). Graph shows the mean, individual data points, and standard deviations of *kipI* mRNA levels relative to WT in BHIS alone for a minimum of three independent experiments. Data were analyzed by one-way ANOVA with Dunnett’s multiple comparison test comparing treated to untreated samples. \**P* <0.01, \*\*\**P* <0.001.

Previous studies demonstrated that some bile salts repress *C. difficile* growth and spore formation through undetermined mechanisms (Sorg and Sonenshein, 2009; Theriot et al., 2016; Usui et al., 2020). Examination of growth in the various bile salts in our hands showed that CDCA, DCA, LCA, GCDCA, and GCA repressed growth, similar to previous findings (**Figure S3**). Based on the induction of *kip* expression and the growth impacts of these bile salts, we chose a representative bile acid, CDCA, to test for potential effects on sporulation through the Kip locus (**Figure S4**). The addition of CDCA decreased spore production in the wild-type and *kip* mutant strains proportionally, with similar decreases in the sporulation ratios across strains. Thus, even with induction of *kip* expression by CDCA, sporulation was not differentially impacted in the wild-type or *kip* mutants, further supporting that KipI and KipA do not have the role in *C. difficile* sporulation regulation observed in *B. subtilis*.

### *C. difficile* uses the toxic byproduct 5-oxoproline (OP) as a nutrient source

As mentioned, the KipOIA oxoprolinase complex detoxifies the metabolic byproduct 5-oxoproline to glutamic acid in both Gram-positive and Gram-negative bacteria (Niehaus et al., 2017; Seddon et al., 1984). In other bacteria, Kip oxoprolinase activity has the dual effect of preventing the accumulation of toxic amounts of OP, while also allowing OP use as a nitrogen source (Niehaus et al., 2017; Ratcliffe et al., 1983). To determine if the predicted *C. difficile* KipOTIA orthologs are involved in 5-oxoproline metabolism, we grew wild-type, the Δ*kipIA* and Δ*kipOTIA* mutants, and complement strains in Defined Minimal Media (DMM) with and without 30 mM 5-oxoproline or 30 mM glutamic acid (**Figure 3, Figure S5**). In DMM, the Δ*kipIA*, and Δ*kipOTIA* mutants grew more slowly than the parent strain, suggesting an intrinsic growth defect in the absence of the Kip complex (**Figure 3D**). The growth defect of the *kip* mutants was partially restored in the presence of glutamic acid (**Figure 3 B,C,F**), suggesting that additional nutrients help to alleviate the growth delay. However, when grown in the presence of OP, the Δ*kipIA* or Δ*kipOTIA* mutants demonstrated further growth retardation than observed in DMM alone, indicating that the Kip complex is both necessary for the utilization of OP and the prevention of deleterious effects by OP. In contrast, the parent strain 630Δ*erm* experienced markedly improved growth and culture density when OP was added to the medium (**Figure 3 A,E**). Remarkably, the growth of the wild-type strain in OP was more robust than the growth observed with the addition of glutamic acid, suggesting that OP is more readily utilized as a nutrient than the metabolic product, glutamic acid. These data demonstrate that the *kipOTIA* operon encodes proteins responsible for 5-oxoprolinase function, and that utilization of 5-oxoproline promotes *C. difficile* growth.

**Figure 3.**
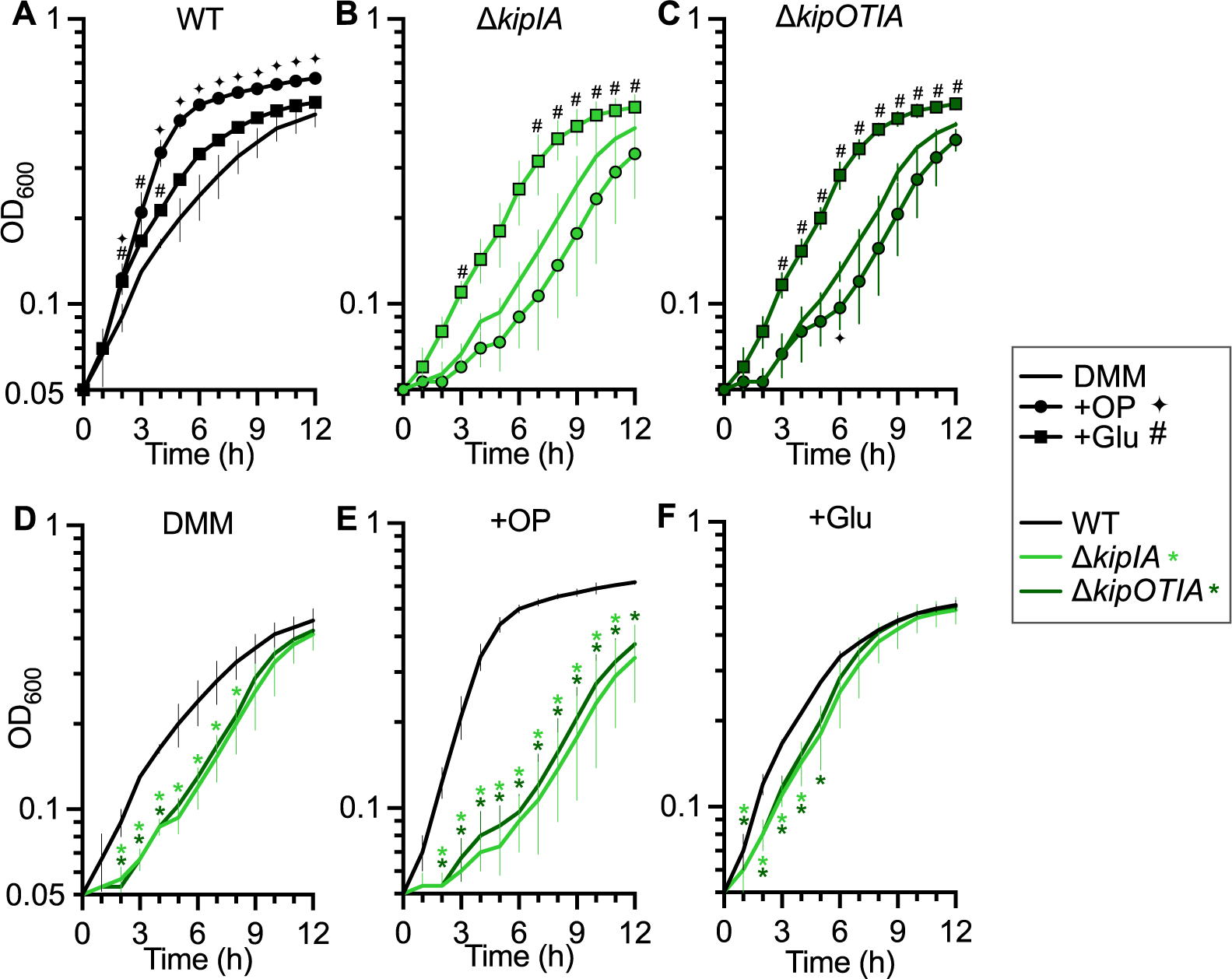
*C. difficile* KipOTIA supports the utilization of 5-oxoproline as a carbon source. **A-C)** Growth of WT (630Δ*erm*), Δ*kipIA* (MC1903), and Δ*kipOTIA* (MC2375) in Defined Minimal Medium (DMM) with and without 30 mM 5-oxoproline (OP) or glutamic acid. **D-F)** Condition-centered display of growth for WT, Δ*kipIA*, and Δ*kipOTIA* in **D)** DMM, **E)** DMM with 30 mM of neutralized 5-oxoproline, and **F)** DMM with 30 mM glutamic acid. The means and SD of three independent experiments are shown. Data in A-C were analyzed by one-way ANOVA with Dunnett’s multiple comparison test at each timepoint to growth in DMM. Data in D-F were analyzed by one-way ANOVA with Dunnett’s multiple comparison test at each timepoint to WT; *, ✦, # *P* <0.05.

Considering that the bile acid CDCA increased the expression of the *kip* genes, we examined whether CDCA could further improve *C. difficile* growth in OP. To do so, we added 0.1 mM CDCA with and without 30 mM OP to the DMM cultures and evaluated the effects of each variable on growth (**Figure 4**). No further enhancement of OP-dependent growth was observed when CDCA and OP were both added to the medium. These data suggest that the addition of CDCA does not enhance the ability of *C. difficile* to utilize 5-oxoproline for growth.

**Figure 4.**
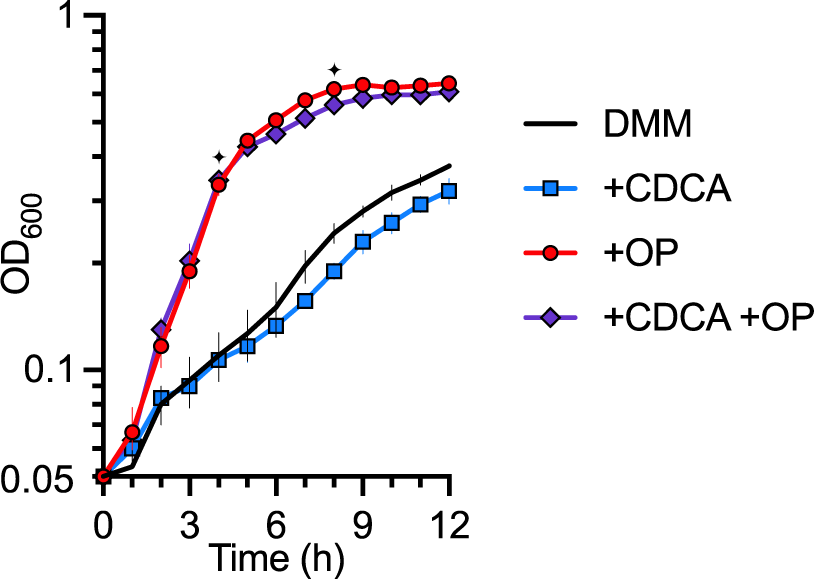
The bile acid CDCA does not enhance *C. difficile* growth in 5-oxoproline. Growth of 630Δ*erm* in Defined Minimal Medium (DMM) with and without 30 mM 5-oxoproline (OP) and/or 0.1 mM of chenodeoxycholate (CDCA). The means and SD of three biological replicates are shown. Data were analyzed by one-way ANOVA with Šídák’s multiple comparison test for each timepoint comparing growth in DMM to +CDCA (*, *P*<0.05) or +OP to +CDCA+OP (✦, *P* <0.05).

As OP has substantial impacts on *C. difficile* growth, we considered that OP utilization may affect sporulation. We extended our sporulation assays to test the impact of exogenous OP on the sporulation frequencies of the *kip* mutants and parent strain, but observed no effect on sporulation outcomes relative to sporulation medium lacking OP (**Figure S4**). Further, we examined the expression of *kipI* in the presence of OP and glutamic acid, and observed only modest changes in *kipI* expression, suggesting that these nutrients do not substantially alter expression of the *kip* operon (**Figure S6**). Thus, OP promotes the growth of *C. difficile* through utilization by the Kip oxoprolinase, but OP does not affect *kip* expression or spore formation.

### The *B. subtilis* oxoprolinase enables the use of OP as a nitrogen source, but not a carbon source

In the initial characterization of the *B. subtilis* Kip/Pxp oxoprolinase, it was observed that the oxoprolinase activity facilitated OP utilization as a nitrogen source (Niehaus et al., 2017). However, the OP-dependent enhancement of growth observed with *C. difficile* suggests that the Kip oxoprolinase allows OP to be used as both a carbon and nitrogen source to support growth. These results led us to question whether the Kip oxoprolinases of *C. difficile* and *B. subtilis* share the capacity for growth enhancement by OP and if *B. subtilis* Kips facilitate the use of OP as a carbon source. To test these hypotheses, we obtained previously generated *kipI* and *kipA* deletion mutants and further investigated their growth profiles in the presence of OP with and without the addition of succinate as a carbon source (Koo et al., 2017). Succinate was chosen as the carbon source to avoid the production of glutamic acid and/or 5-oxoproline. The growth of wild-type *B. subtilis, ΔkipI,* and *ΔkipA* mutants were compared in Niehaus Defined Medium (NDM) with added succinate, OP, or both (**Figure 5**). When succinate served as the sole carbon source, poor growth was observed for all strains, but the *ΔkipI* and *ΔkipA* mutants demonstrated significantly less growth than the parent strain (**Figure 5A**). Importantly, OP alone did not support the growth of any *B. subtilis* strain and led to decreased cell density for both *kip* mutants (**Figure 5B**). Addition of both succinate and OP to the medium resulted in considerable growth for the parent strain that was not observed for either the Δ*kipI* or Δ*kipA* mutant (**Figure 5C**). These observations reinforce the previous findings of OP toxicity in the absence of the *kip* genes and demonstrate that OP is not an adequate sole carbon source for *B. subtilis*.

**Figure 5.**
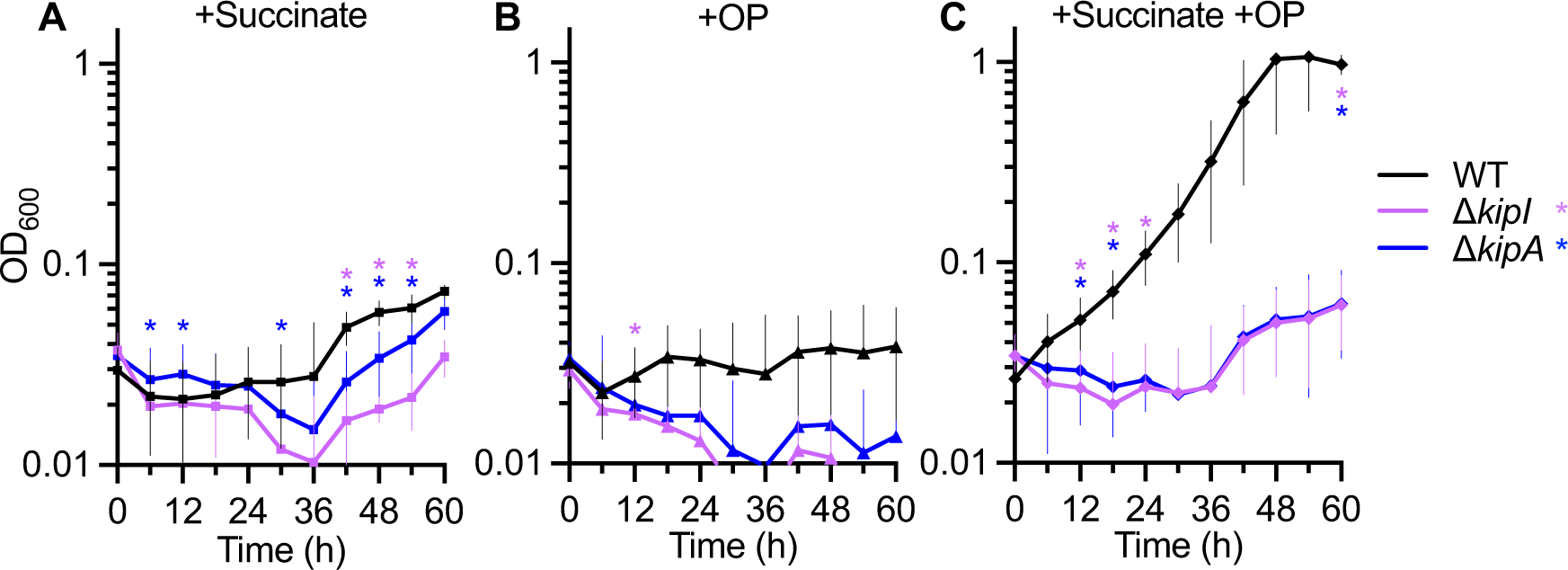
The *B. subtilis* Kip oxoprolinase enables the use of 5-oxoproline as a nitrogen source, but not a carbon source. **A-C)** Growth of *B. subtilis* 1A1 (wild-type), Δ*kipI* (MC2624), and Δ*kipA* (MC2625) in Niehaus Defined Medium (NDM) with or without 30 mM sodium succinate (SS), 30 mM 5-oxoproline (OP), or 30 mM OP and 30 mM SS. The means and SD of three independent experiments are shown. Data were analyzed by one-way ANOVA with Dunnett’s multiple comparison test at each timepoint comparing mutants to wild-type; *, *, *P* <0.05.

## DISCUSSION

In this study, we investigated the role of the KipOTIA factors in the regulation of sporulation and the ability of *C. difficile* to detoxify the metabolite 5-oxoproline. We found that KipI and KipA do not have major effects on *C. difficile* sporulation, in contrast to *B. subtilis*, in which KipI impedes sporulation (Wang et al., 1997). The absence of a sporulation effect by the *C. difficile* Kip proteins is not completely surprising given the considerable differences in the genetic pathways and factors involved in sporulation initiation between *B. subtilis* and *C. difficile* (Edwards et al., 2022; Lee et al., 2022; Mehdizadeh Gohari et al., 2024). In *B. subtilis,* KipI binds to the sporulation activating kinase, KinA to repress spore formation (Jacques et al., 2011, 2008). However, *C. difficile* does not encode a clear KinA ortholog and the phosphotransfer proteins that function in the regulation of sporulation initiation appear to suppress spore formation (Edwards et al., 2022). Moreover, it is not known how the *Bacillus* KinA-KipI-KipA interactions are controlled or if such signals are present or relevant for *C. difficile.* While *C. difficile* Kips have no apparent role in sporulation, there are many sporulating species that encode Kip orthologs, but their functions have not been investigated. Hence, it is possible that other Clostridial KipI proteins function as sporulation kinase inhibitors—most likely those with histidine kinases that activate Spo0A.

The most impactful result of this investigation was the finding that the KipOTIA complex not only functions as a 5-oxoprolinase, but that 5-oxoproline supports robust *C. difficile* growth. For *B. subtilis,* the conversion of OP to glutamic acid provides a detoxification function and supplies an excellent nitrogen source for the bacterium, but glutamic acid cannot serve as a sole carbon source for *B. subtilis* (Belitsky et al., 2000; Commichau et al., 2008). The robust growth observed for wild-type *C. difficile* with OP suggests that KipOTIA facilitates the use of OP as both a carbon and a nitrogen source (**Fig. 3**). However, how KipOTIA metabolizes OP to generate energy is not clear. We observed greater cell growth of *C. difficile* in OP than in equal concentrations of glutamic acid, which is curious. The conversion of OP to glutamic acid costs energy, so cells would be expected to grow better with glutamic acid than with OP. The growth differences for *C. difficile* in OP and glutamic acid suggest that OP can be utilized by mechanisms that glutamic acid cannot.

It was previously observed by Fletcher et al. that the concentration of OP increases during *C. difficile* colonization in a mouse model of infection, though the cause of the OP increase is not known (Fletcher et al., 2018). The presence of both exogenous and endogenously generated OP necessitates detoxification by both the intestinal microbiota and the host. The fact that *C. difficile* growth is enhanced by a metabolite that is toxic to other species could greatly impact competition with other intestinal microbes and may provide a significant nutritional advantage to support *C. difficile* proliferation during infection. Further, the growth advantages afforded to *C. difficile* by KipOTIA-mediated 5-oxoproline metabolism may be shared by other species that encode Kip orthologs.

## MATERIALS AND METHODS

### Strain and plasmid construction

See **Table S1** for bacterial strains and plasmids used in this study. The 630 strain genome (GenBank accession no. NC_009089.1) was used to design the primers for *C. difficile* listed in **Table S2**. *C. difficile* 630Δ*erm* genomic DNA, pRT1099, and pJIR1457 were used as templates for PCR reactions and mutant construction. Vector construction details are provided in **Table S3**. All *C. difficile* mutants were made using the pseudo-suicide allele-coupled exchange, as previously described (Edwards et al., 2022; Peltier et al., 2020) All vectors were verified by whole plasmid sequencing using Oxford Nanopore Technology (Plasmidsaurus). *C. difficile* strains were confirmed by PCR analysis and whole-genome sequencing. The *B. subtilis* Δ*kipI* and Δ*kipA* mutants were created by natural competence transformation of strain 1A1 with genomic DNA from strains BKE04080 and BKE04090 respectively (Koo et al., 2017).

### Bacterial growth assays and conditions

All *C. difficile* strains were cultured in a Coy anaerobic chamber at 37°C with an atmosphere of 10% H2, 5% CO2, and balanced with 85% N2 as previously described (Edwards et al., 2013; Smith et al., 1981). *C. difficile* strains were grown in brain-heart infusion medium supplemented with yeast extract (Bacto) (Putnam et al., 2013; Sorg and Dineen, 2009). For plasmid maintenance and integrant selection in *C. difficile,* cultures were supplemented with 2-10 µg/mL of thiamphenicol (Sigma-Aldrich) or 500 µg/mL spectinomycin (Thermo Scientific). Counterselection against *E. coli* post-conjugation was achieved using 100 µg/mL of kanamycin. For growth curves and spore assays, strains were grown overnight in BHIS with 0.1% taurocholate (TA; Sigma-Aldrich) and 0.2% fructose (D-fructose, Fisher Chemical) added as needed to induce *C. difficile* germination and prevent sporulation (Putnam et al., 2013; Sorg and Dineen, 2009). 5-oxoproline was neutralized and filter sterilized immediately before use. All bile salts were diluted in dH2O except for lithocholate, which was solubilized in 95% ethanol.

*Escherichia coli* strains were grown aerobically at 37°C in LB medium (Lennox) supplemented with 20 µg/mL chloramphenicol, 100 µg/mL spectinomycin, or 100 µg/mL ampicillin (Sigma-Aldrich) as needed for plasmid maintenance.

For *C. difficile* growth curves, strains were grown in a chemically defined minimal medium (DMM), loosely based on modified CDMM (Karasawa et al., 1995; Rizvi et al., 2023); (**Table S4**). Neutralized 5-oxoproline and/or glutamic acid were added to DMM as indicated. Each medium was made fresh and used within 12 h. Cultures were inoculated in BHIS with 0.1% TA and 0.2% fructose, and grown to turbidity. Active cultures were diluted with BHIS, grown to an OD600=0.5, pelleted for 10 min at 3214 x *g*, resuspended to an OD600=0.5 in DMM and 5 ml was inoculated into 45 ml of DMM.

For *B. subtilis* growth curves, cultures were grown to logarithmic phase in LB broth (Lennox), pelleted for 10 min at 3214 x *g*, and washed twice with 5 mL modified Niehaus Defined Media (NDM, **Table S5**) without a carbon or nitrogen source (Niehaus et al., 2017). Cultures were resuspended in NDM to an OD600=0.5 and diluted 1:10 into 1.8 mL of NDM supplemented with 30 mM sodium succinate, 30 mM 5-oxoproline, or 30 mM glutamic acid, as indicated. Cultures were grown in a 24-well suspension cell plate (Sarstedt) for 72 h at 37°C with continuous double orbital shaking at 548 cpm in a BioTek Synergy H1 microplate reader. The OD600 was measured for each culture at 30 minute intervals and visualized using the Gen5 software version 3.11.19. Growth data were analyzed by one-way ANOVA with Dunnett’s multiple comparison test or Student’s two-tailed *t*-tests as indicated.

### Whole Genome Sequencing

*C. difficile* strains were grown in BHIS, genomic DNA extracted and prepared for whole genome sequencing as previously detailed (A. N. Edwards et al., 2016; Edwards and McBride, 2023; Harju et al., 2004). Library preparation and Illumina sequencing was performed by SeqCenter (Pittsburgh, PA) as previously described (Edwards and McBride, 2023). Demultiplexing, quality control, and adapter trimming was performed with bcl-convert (v3.9.3 for MC2375, v4.1.5 for MC2519 and MC2520; https://support-docs.illumina.com/SW/BCL_Convert/Content/SW/FrontPages/BCL_Convert.htm). Geneious Prime (v2023.1.2 and v2021.2.2) was used to trim reads using the BBDuk plug-in and then mapped to the *C. difficile* reference genome NC_009089. The Bowtie2 plugin was used to find SNPs or InDels under default settings with a minimum variant frequency of 0.9 (Langmead and Salzberg, 2012). Genome sequence files were deposited into NCBI Sequence Read Archive (SRA) BioProject PRJNA1085672.

### Sporulation assays

Ethanol-resistant sporulation assays were performed as previously described (Childress et al., 2016; DiCandia et al., 2022; Edwards et al., 2022; Edwards and McBride, 2017, 2016). Briefly, *C. difficile* strains were grown in BHIS broth supplemented with 0.1% TA and 0.2% fructose to germinate spores and prevent spore formation, then diluted in BHIS and grown to log-phase (OD600 =0.5). Cultures were plated onto 70:30 sporulation agar, with or without 0.5 mM chenodeoxycholate (Calbiochem), or 30 mM of neutralized 5-oxoproline (L-pyroglutamic acid, Sigma-Aldrich), as indicated. After 24 h, cells were plated for vegetative cell counts or treated with ethanol, diluted, and plated for spore enumeration on BHIS with 0.1% TA. Sporulation frequency was calculated as the number of ethanol-resistant spores divided by the total number of cells. A *spo0A* mutant (MC310) was used as a negative control to verify vegetative cell death during ethanol treatment. At least four independent experiments were performed, and sporulation frequency data were analyzed by one-way or two-way analysis of variance (ANOVA) with Dunnett’s or Šídák’s multiple comparison test, as indicated (GraphPad Prism v10.0.2).

### Quantitative reverse transcription PCR analysis (qRT-PCR)

*C. difficile* was grown on 70:30 agar or in BHIS broth, as indicated, and samples processed for RNA extraction. Culture samples were taken at H12 (12 h after cultures were applied to the plate) for 70:30 agar samples or at an OD600=0.5 for BHIS growth curve samples. Cultures were resuspended in ethanol:acetone:dH2O solution and stored at - 80°C. RNA was isolated, Dnase-I treated (Ambion), and cDNA was synthesized (Bioline) using random hexamers as previously described (Dineen et al., 2010; Edwards et al., 2014; Suarez et al., 2013). qRT-PCR analysis was performed on 50 ng of cDNA using the SensiFAST SYBR & Fluorescein kit (Bioline) with a Roche Lightcycler 96. qRT-PCR primers are listed in **Table S2** and each qRT-PCR reaction was performed in technical triplicate with at least three biological replicates. The ΔΔCt method was used to analyze results and normalize expression to the internal control transcript *rpoC* for relative quantification (Schmittgen and Livak, 2008). Statistical analysis of qRT-PCR data was performed using a one-way ANOVA as indicated (GraphPad Prism v10.0.2).

### RNA-Seq

*C. difficile* strains 630Δ*erm* pMC123 (MC324) and 630Δ*erm* Δ*kipIA* pMC123 (MC1970) were grown in 70:30 broth (pH=8.0) with 2 µg/mL thiamphenicol to an OD600=0.5 and samples were taken as described above for qRT-PCR. RNA was isolated and treated with Dnase-I, and samples were sent to Seq-Center for sequencing. Illumina’s Stranded Total RNA Prep Ligation with Ribo-Zero Plus kit and 10bp IDK for Illumina indices was used for library preparation. Sequencing was performed on a NextSeq2000 giving 2×50bp reads. Demultiplexing, quality control, and adapter trimming was performed with bcl-convert (v3.9.3; see reference above for whole genome sequencing). Geneious Prime v2022.2.2 was used to map reads to the 630Δ*erm* reference genome (NC_009089.1) and expression levels were calculated and compared using DESeq2 (Love et al., 2014). DESeq2 performs the Wald test to calculate *P* values which are adjusted by the Benjamini-Hochberg test (Love et al., 2014). Raw RNA sequencing reads were deposited to the NCBI Sequence Read Archive (SRA) BioProject PRJNA1085672.

## ACKNOWLEDGEMENTS

The authors would like to thank the members of the McBride lab for their helpful feedback and suggestions during the completion of this manuscript. We would also like to thank Sarah Anderson for performing preliminary work characterizing the response of *C. difficile* to deoxycholate. This research was supported by the U.S. National Institutes of Health through research grants AI116933 and AI156052 to S.M.M. and AI106699 to C.D.L. The content of this manuscript is solely the responsibility of the authors and does not necessarily reflect the official views of the National Institutes of Health.

## REFERENCES

Ali, S., Moore, G., Wilson, A.P., 2011. Spread and persistence of Clostridium difficile spores during and after cleaning with sporicidal disinfectants. The Journal of hospital infection 79, 97–8. 10.1016/j.jhin.2011.06.010

Antunes, A., Camiade, E., Monot, M., Courtois, E., Barbut, F., Sernova, N.V., Rodionov, D.A., Martin-Verstraete, I., Dupuy, B., 2012. Global transcriptional control by glucose and carbon regulator CcpA in Clostridium difficile. Nucleic acids research 40, 10701–18. 10.1093/nar/gks864

Awadé, A., Cleuziat, Ph., Gonzalès, Th., Robert-Baudouy, J., 1992. Characterization of the pcp gene encoding the pyrrolidone carboxyl peptidase of Bacillus subtilis. FEBS Letters 305, 67–73. 10.1016/0014-5793(92)80656-2

Awadé, A.C., Cleuziat, Ph., GonzalèS, Th., Robert-Baudouy, J., 1994. Pyrrolidone carboxyl peptidase (Pcp): An enzyme that removes pyroglutamic acid (pGlu) from pGlu-peptides and pGlu-proteins. Proteins: Structure, Function, and Bioinformatics 20, 34–51. 10.1002/prot.340200106

Belitsky, B.R., Wray, L.V., Fisher, S.H., Bohannon, D.E., Sonenshein, A.L., 2000. Role of TnrA in nitrogen source-dependent repression of Bacillus subtilis glutamate synthase gene expression. Journal of bacteriology 182, 5939–47.

Bouillaut, L., Dubois, T., Francis, M.B., Daou, N., Monot, M., Sorg, J.A., Sonenshein, A.L., Dupuy, B., 2019. Role of the global regulator Rex in control of NAD ^+^ - regeneration in *Clostridioides (Clostridium) difficile*. Mol Microbiol 111, 1671– 1688. 10.1111/mmi.14245

Childress, K.O., Edwards, A.N., Nawrocki, K.L., Woods, E.C., Anderson, S.E., McBride, S.M., 2016. The Phosphotransfer Protein CD1492 Represses Sporulation Initiation in Clostridium difficile. Infection and immunity. 10.1128/IAI.00735-16

Commichau, F.M., Gunka, K., Landmann, J.J., Stülke, J., 2008. Glutamate Metabolism in *Bacillus subtilis* : Gene Expression and Enzyme Activities Evolved To Avoid Futile Cycles and To Allow Rapid Responses to Perturbations of the System. J Bacteriol 190, 3557–3564. 10.1128/JB.00099-08

DiCandia, M.A., Edwards, A.N., Jones, J.B., Swaim, G.L., Mills, B.D., McBride, S.M., 2022. Identification of Functional Spo0A Residues Critical for Sporulation in Clostridioides difficile. Journal of Molecular Biology 434, 167641. 10.1016/j.jmb.2022.167641

Dineen, S.S., McBride, S.M., Sonenshein, A.L., 2010. Integration of metabolism and virulence by Clostridium difficile CodY. Journal of bacteriology 192, 5350–62. 10.1128/JB.00341-10

Doolittle, R.F., Armentrout, R.W., 1968. Pyrrolidonyl peptidase. Enzyme for selective removal of pyrrolidonecarboxylic acid residues from polypeptides. Biochemistry 7, 516–521. 10.1021/bi00842a005

Dubberke, E.R., Olsen, M.A., 2012. Burden of Clostridium difficile on the healthcare system. Clinical infectious diseases : an official publication of the Infectious Diseases Society of America 55 Suppl 2, S88–92. 10.1093/cid/cis335

Edwards, A.N., Karim, S.T., Pascual, R.A., Jowhar, L.M., Anderson, S.E., McBride, S.M., 2016. Chemical and Stress Resistances of Clostridium difficile Spores and Vegetative Cells. Frontiers in microbiology 7, 1698.

Edwards, A.N., McBride, S.M., 2023. The RgaS-RgaR two-component system promotes Clostridioides difficile sporulation through a small RNA and the Agr1 system. PLoS Genet 19, e1010841. 10.1371/journal.pgen.1010841

Edwards, A.N., McBride, S.M., 2017. Determination of the in vitro Sporulation Frequency of Clostridium difficile. Bio-protocol 7. 10.21769/BioProtoc.2125

Edwards, A.N., McBride, S.M., 2016. Isolating and Purifying Clostridium difficile Spores. Methods Mol Biol 1476, 117–28. 10.1007/978-1-4939-6361-4_9

Edwards, A.N., Nawrocki, K.L., McBride, S.M., 2014. Conserved oligopeptide permeases modulate sporulation initiation in *Clostridium difficile*. Infection and immunity 82, 4276–91. 10.1128/IAI.02323-14

Edwards, A.N., Suarez, J.M., McBride, S.M., 2013. Culturing and maintaining *Clostridium difficile* in an anaerobic environment. Journal of visualized experiments : JoVE e50787. 10.3791/50787

Edwards, A. N., Tamayo, R., McBride, S.M., 2016. A novel regulator controls Clostridium difficile sporulation, motility and toxin production. Molecular microbiology 100, 954–71. 10.1111/mmi.13361

Edwards, A.N., Wetzel, D., DiCandia, M.A., McBride, S.M., 2022. Three Orphan Histidine Kinases Inhibit Clostridioides difficile Sporulation. J Bacteriol 204, e0010622. 10.1128/jb.00106-22

Errington, J., 2003. Regulation of endospore formation in Bacillus subtilis. Nature reviews. Microbiology 1, 117–26. 10.1038/nrmicro750

Finn, E., Andersson, F.L., Madin-Warburton, M., 2021. Burden of Clostridioides difficile infection (CDI) - a systematic review of the epidemiology of primary and recurrent CDI. BMC Infect Dis 21, 456. 10.1186/s12879-021-06147-y

Fletcher, J.R., Erwin, S., Lanzas, C., Theriot, C.M., 2018. Shifts in the Gut Metabolome and Clostridium difficile Transcriptome throughout Colonization and Infection in a Mouse Model. mSphere 3. 10.1128/mSphere.00089-18

Harju, S., Fedosyuk, H., Peterson, K.R., 2004. Rapid isolation of yeast genomic DNA: Bust n’ Grab. BMC biotechnology 4, 8. 10.1186/1472-6750-4-8

Jacques, D.A., Langley, D.B., Hynson, R.M., Whitten, A.E., Kwan, A., Guss, J.M., Trewhella, J., 2011. A novel structure of an antikinase and its inhibitor. Journal of molecular biology 405, 214–26. 10.1016/j.jmb.2010.10.047

Jacques, D.A., Langley, D.B., Jeffries, C.M., Cunningham, K.A., Burkholder, W.F., Guss, J.M., Trewhella, J., 2008. Histidine Kinase Regulation by a Cyclophilin-like Inhibitor. Journal of Molecular Biology 384, 422–435. 10.1016/j.jmb.2008.09.017

Karasawa, T., Ikoma, S., Yamakawa, K., Nakamura, S., 1995. A defined growth medium for Clostridium difficile. Microbiology 141, 371–375. 10.1099/13500872-141-2-371

Koo, B.-M., Kritikos, G., Farelli, J.D., Todor, H., Tong, K., Kimsey, H., Wapinski, I., Galardini, M., Cabal, A., Peters, J.M., Hachmann, A.-B., Rudner, D.Z., Allen, K.N., Typas, A., Gross, C.A., 2017. Construction and Analysis of Two Genome-Scale Deletion Libraries for Bacillus subtilis. Cell Systems 4, 291–305.e7. 10.1016/j.cels.2016.12.013

Kumar, A., Bachhawat, A.K., 2010. OXP1/YKL215c encodes an ATP-dependent 5-oxoprolinase in Saccharomyces cerevisiae: functional characterization, domain structure and identification of actin-like ATP-binding motifs in eukaryotic 5-oxoprolinases: OXP1 encodes a functional 5-oxoprolinase in S. cerevisiae. FEMS Yeast Research 10, 394–401. 10.1111/j.1567-1364.2010.00619.x

Langmead, B., Salzberg, S.L., 2012. Fast gapped-read alignment with Bowtie 2. Nat Methods 9, 357–359. 10.1038/nmeth.1923

Lee, C.D., Rizvi, A., Edwards, A.N., DiCandia, M.A., Vargas Cuebas, G.G., Monteiro, M.P., McBride, S.M., 2022. Genetic mechanisms governing sporulation initiation in Clostridioides difficile. Current Opinion in Microbiology 66, 32–38. 10.1016/j.mib.2021.12.001

Li, L.Y., Seddon, A.P., Meister, A., 1988. Interaction of the protein components of 5-oxoprolinase. Substrate-dependent enzyme complex formation. Journal of Biological Chemistry 263, 6495–6501. 10.1016/S0021-9258(18)68670-3

Love, M.I., Huber, W., Anders, S., 2014. Moderated estimation of fold change and dispersion for RNA-seq data with DESeq2. Genome Biology 15, 550. 10.1186/s13059-014-0550-8

Mehdizadeh Gohari, I., Edwards, A.N., McBride, S.M., McClane, B.A., 2024. The impact of orphan histidine kinases and phosphotransfer proteins on the regulation of clostridial sporulation initiation. mBio 15, e02248–23. 10.1128/mbio.02248-23

Neumann-Schaal, M., Jahn, D., Schmidt-Hohagen, K., 2019. Metabolism the Difficile Way: The Key to the Success of the Pathogen Clostridioides difficile. Frontiers in microbiology 10, 219. 10.3389/fmicb.2019.00219

Niehaus, T.D., Elbadawi-Sidhu, M., De Crécy-Lagard, V., Fiehn, O., Hanson, A.D., 2017. Discovery of a widespread prokaryotic 5-oxoprolinase that was hiding in plain sight. Journal of Biological Chemistry 292, 16360–16367. 10.1074/jbc.M117.805028

Northfield, T.C., McColl, I., 1973. Postprandial concentrations of free and conjugated bile acids down the length of the normal human small intestine. Gut 14, 513–518. 10.1136/gut.14.7.513

Park, C.B., Lee, S.B., Ryu, D.D.Y., 2001. l-Pyroglutamate Spontaneously Formed from l-Glutamate Inhibits Growth of the Hyperthermophilic Archaeon Sulfolobus solfataricus. Appl Environ Microbiol 67, 3650–3654. 10.1128/AEM.67.8.3650-3654.2001

Peltier, J., Hamiot, A., Garneau, J.R., Boudry, P., Maikova, A., Hajnsdorf, E., Fortier, L.-C., Dupuy, B., Soutourina, O., 2020. Type I toxin-antitoxin systems contribute to the maintenance of mobile genetic elements in Clostridioides difficile. Commun Biol 3, 718. 10.1038/s42003-020-01448-5

Putnam, E.E., Nock, A.M., Lawley, T.D., Shen, A., 2013. SpoIVA and SipL are Clostridium difficile spore morphogenetic proteins. Journal of bacteriology 195, 1214–25. 10.1128/JB.02181-12

Ratcliffe, H.D., Drozd, J.W., Bull, A.T., 1983. The Utilization of 5-Oxoproline, Ammonia and Glutamine by Rhizobium leguminosarum in Chemostat Culture. Microbiology. 10.1099/00221287-129-6-1707

Rizvi, A., Vargas-Cuebas, G., Edwards, A.N., DiCandia, M.A., Carter, Z.A., Lee, C.D., Monteiro, M.P., McBride, S.M., 2023. Glycine fermentation by C. difficile promotes virulence and spore formation, and is induced by host cathelicidin. Infection and Immunity 91, e00319–23. 10.1128/iai.00319-23

Schmittgen, T.D., Livak, K.J., 2008. Analyzing real-time PCR data by the comparative C(T) method. Nature protocols 3, 1101–8.

Seddon, A.P., Li, L.Y., Meister, A., 1984. Resolution of 5-oxo-L-prolinase into a 5-oxo-L-proline-dependent ATPase and a coupling protein. Journal of Biological Chemistry 259, 8091–8094. 10.1016/S0021-9258(17)39697-7

Seddon, A.P., Meister, A., 1986. Trapping of an intermediate in the reaction catalyzed by 5-oxoprolinase. Journal of Biological Chemistry 261, 11538–11543. 10.1016/S0021-9258(18)67276-X

Shen, A., 2020. Clostridioides difficile Spore Formation and Germination: New Insights and Opportunities for Intervention. Annu. Rev. Microbiol. 74, 545–566. 10.1146/annurev-micro-011320-011321

Shen, A., Edwards, A.N., Sarker, M.R., Paredes-Sabja, D., 2019. Sporulation and Germination in Clostridial Pathogens. Microbiology spectrum 7. 10.1128/microbiolspec.GPP3-0017-2018

Smith, C.J., Markowitz, S.M., Macrina, F.L., 1981. Transferable tetracycline resistance in Clostridium difficile. Antimicrobial agents and chemotherapy 19, 997–1003.

Sonenshein, A.L., 2007. Control of key metabolic intersections in *Bacillus subtilis*. Nature reviews. Microbiology 5, 917–27. 10.1038/nrmicro1772

Sonenshein, A.L., 2000. Control of sporulation initiation in *Bacillus subtilis*. Current opinion in microbiology 3, 561–6.

Sorg, J.A., Dineen, S.S., 2009. Laboratory maintenance of Clostridium difficile. Current protocols in microbiology Chapter 9, Unit9A 1. 10.1002/9780471729259.mc09a01s12

Sorg, J.A., Sonenshein, A.L., 2009. Chenodeoxycholate is an inhibitor of Clostridium difficile spore germination. Journal of bacteriology 191, 1115–7. 10.1128/JB.01260-08

Suarez, J.M., Edwards, A.N., McBride, S.M., 2013. The *Clostridium difficile cpr* locus is regulated by a noncontiguous two-component system in response to type A and B lantibiotics. Journal of bacteriology 195, 2621–31. 10.1128/JB.00166-13

Theriot, C.M., Bowman, A.A., Young, V.B., 2016. Antibiotic-Induced Alterations of the Gut Microbiota Alter Secondary Bile Acid Production and Allow for Clostridium difficile Spore Germination and Outgrowth in the Large Intestine. mSphere 1. 10.1128/mSphere.00045-15

Usui, Y., Ayibieke, A., Kamiichi, Y., Okugawa, S., Moriya, K., Tohda, S., Saito, R., 2020. Impact of deoxycholate on Clostridioides difficile growth, toxin production, and sporulation. Heliyon 6, e03717. 10.1016/j.heliyon.2020.e03717

Vetter, A.M., Helmecke, J., Schomburg, D., Neumann-Schaal, M., 2019. The Impact of Pyroglutamate: *Sulfolobus acidocaldarius* Has a Growth Advantage over *Saccharolobus solfataricus* in Glutamate-Containing Media. Archaea 2019, e3208051. 10.1155/2019/3208051

Wang, L., Grau, R., Perego, M., Hoch, J.A., 1997. A novel histidine kinase inhibitor regulating development in Bacillus subtilis. Genes and Development 11, 2569– 2579. 10.1101/gad.11.19.2569

