## Supplemental Figures for "KipOTIA detoxifies 5-oxoproline and promotes the growth of *Clostridioides difficile*"

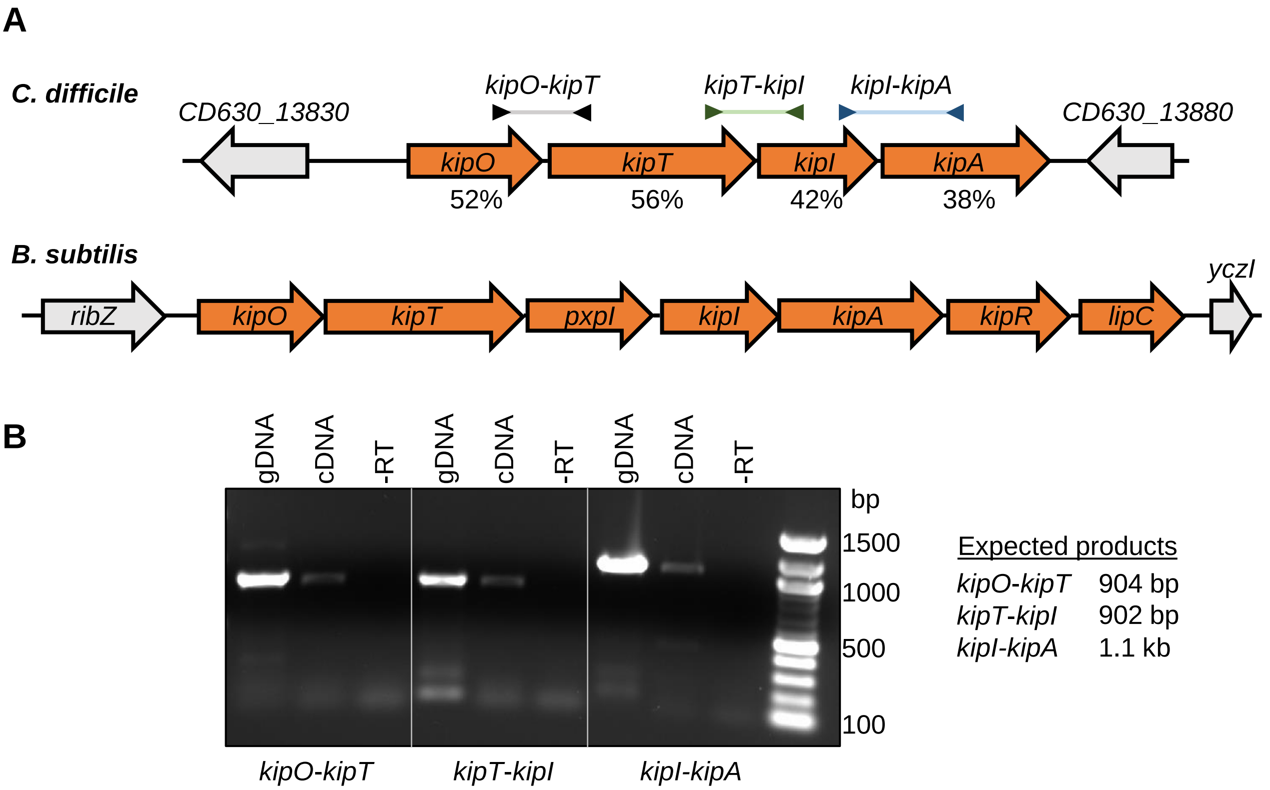


**Figure S1. *C. difficile kipOTIA* is transcribed as an operon. A)** The organization of the *kip* operons in *C. difficile* (*CD630_13840-CD630_13870*) and *B. subtilis* (*BSU04050-BSU04110*). Arrows above the *C. difficile kipOTIA* operon indicate PCR segments shown in B. Numbers below the *C. difficile kipOTIA* operon indicate percent identity to the corresponding proteins in *B. subtilis*. **B)** Strains were grown on 70:30 agar and samples were collected at $H_{12}$ for RNA and cDNA generation as described in the Materials and Methods. Nested PCR was performed using 50 ng genomic DNA (gDNA, positive control), cDNA, or RNA without reverse transcriptase (-RT, negative control). Primer pairs are as follows: *kipO-kipT*, oMC2868/oMC2869; *kipT-kipI*, oMC2870/oMC502; *kipI-kipA*, oMC501/oMC504. PCR products were visualized on a 0.7% agarose gel.

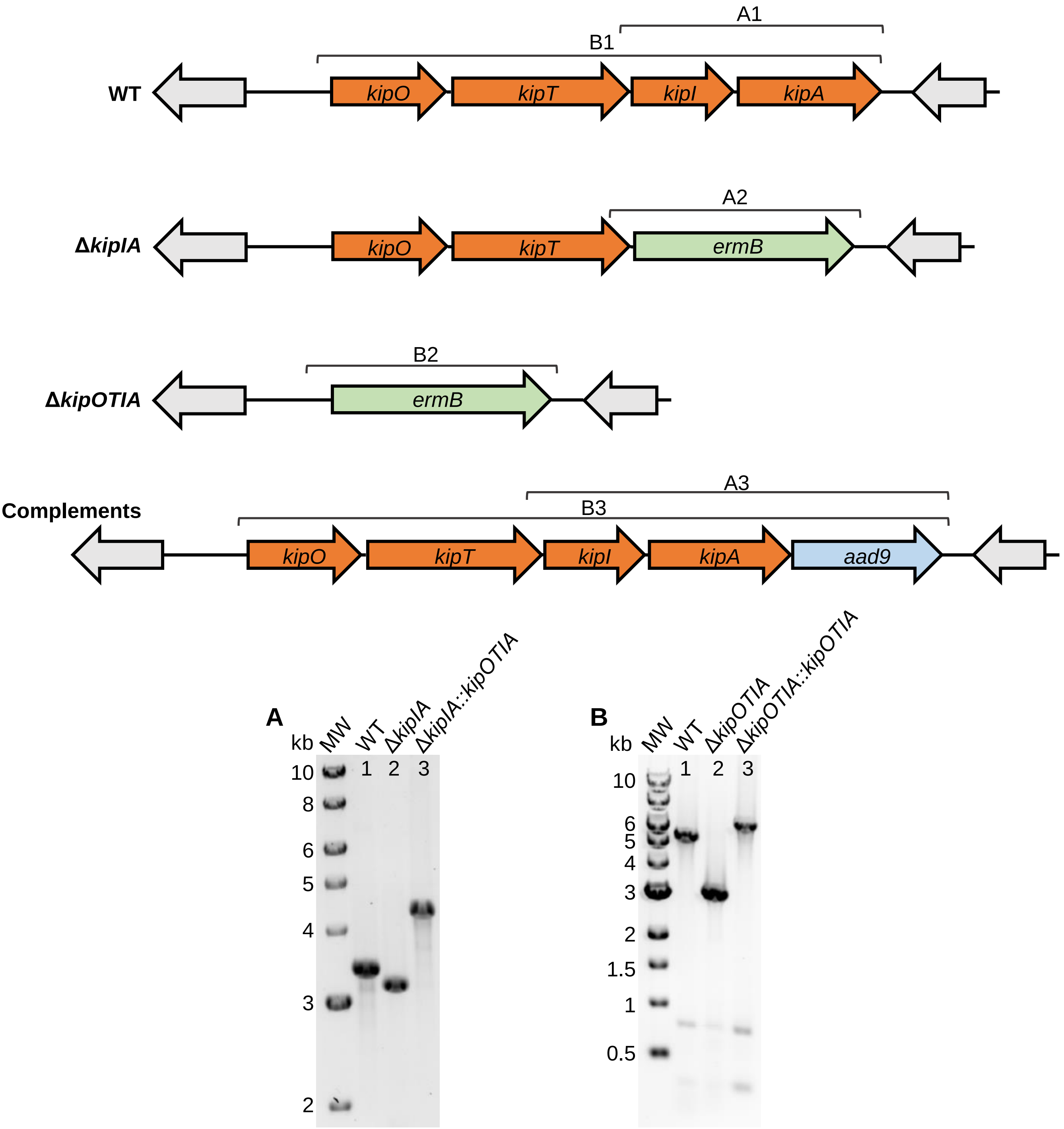


**Figure S2. Construction and confirmation of *C. difficile kip* mutants and complemented strains. A)** PCR confirmation of *kipIA* deletion and complementation with *kipOTIA*::*aad9*. Expected PCR products using primers flanking *kipIA* (oMC2820/2823) are 3,543 bp for WT (630Δ*erm*), 3,354 bp for Δ*kipIA*::*erm* (MC1903), and 4,588 bp for Δ*kipIA*::*kipOTIA*::*aad9* (MC2519). **B)** PCR confirmation of *kipOTIA* deletion and complementation with *kipOTIA*::*aad9.* Expected PCR products using primers flanking *kipOTIA* (oMC3538/2823) are 5,473 bp for WT, 3,224 bp for Δ*kipOTIA*::*erm* (MC2375), and 6,518 bp for Δ*kipOTIA*::*kipOTIA*::*aad9* (MC2520).

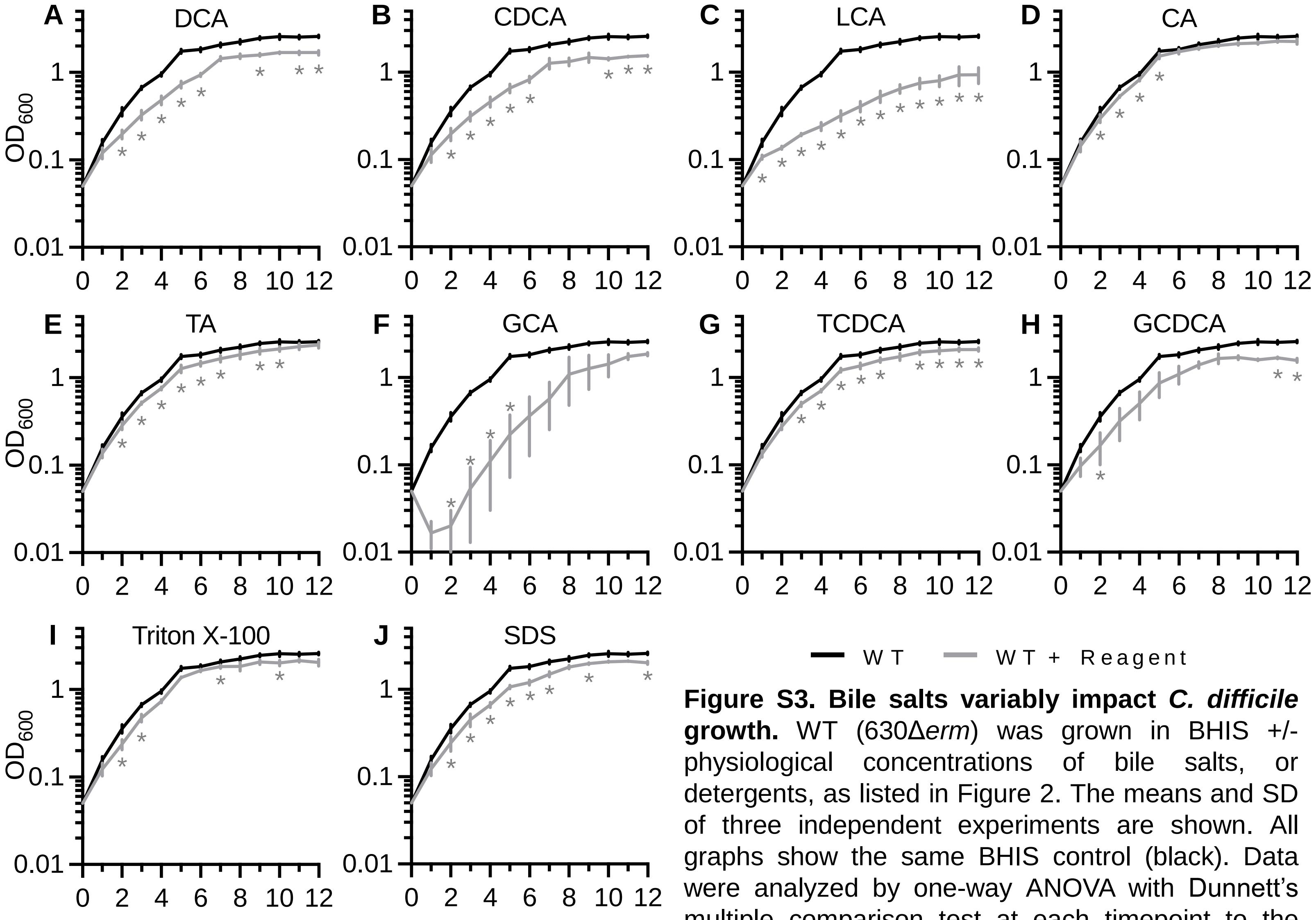


**Figure S3. Bile salts variably impact *C. difficile* growth.** WT (630Δ*erm*) was grown in BHIS +/- physiological concentrations of bile salts, or detergents, as listed in Figure 2. The means and SD of three independent experiments are shown. All graphs show the same BHIS control (black). Data were analyzed by one-way ANOVA with Dunnett's multiple comparison test at each timepoint to the control. **P* <0.05.

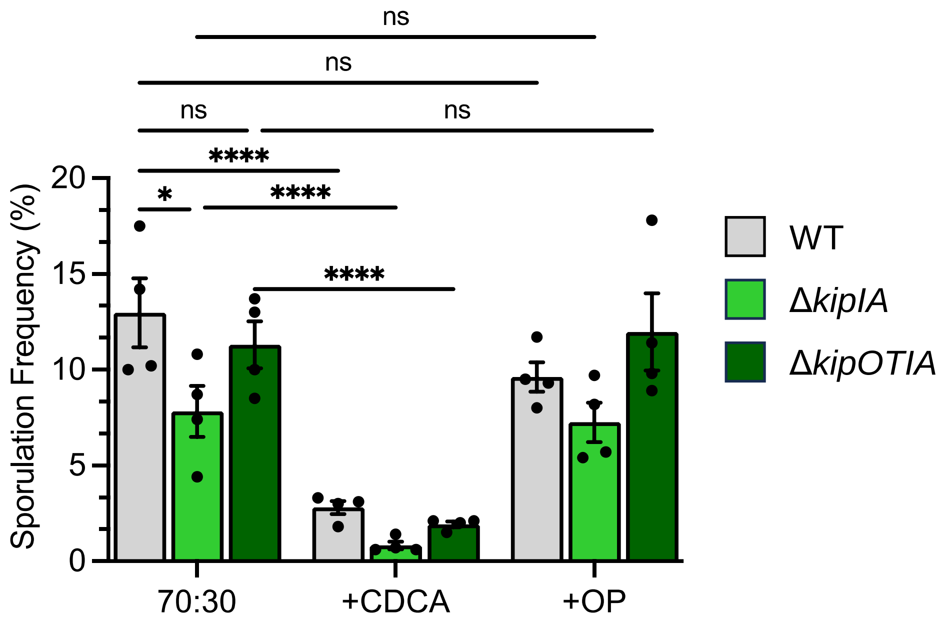


**Figure S4. KipOTIA does not inhibit sporulation under conditions associated with its expression or function.** Ethanol-resistant sporulation frequencies of *C. difficile* strain 630Δ*erm* (WT), Δ*kipIA* mutant (MC1903), and the Δ*kipOTIA* mutant (MC2375) grown for 24 h on 70:30 sporulation agar +/- 0.5 mM chenodeoxycholate (CDCA) or 30 mM of neutralized 5-oxoproline (OP). The means and SEM of four independent experiments are shown. Data were analyzed by two-way ANOVA with Šídák's multiple comparison test; **P*<0.05, ***P*<0.01, *****P*<0.0001.

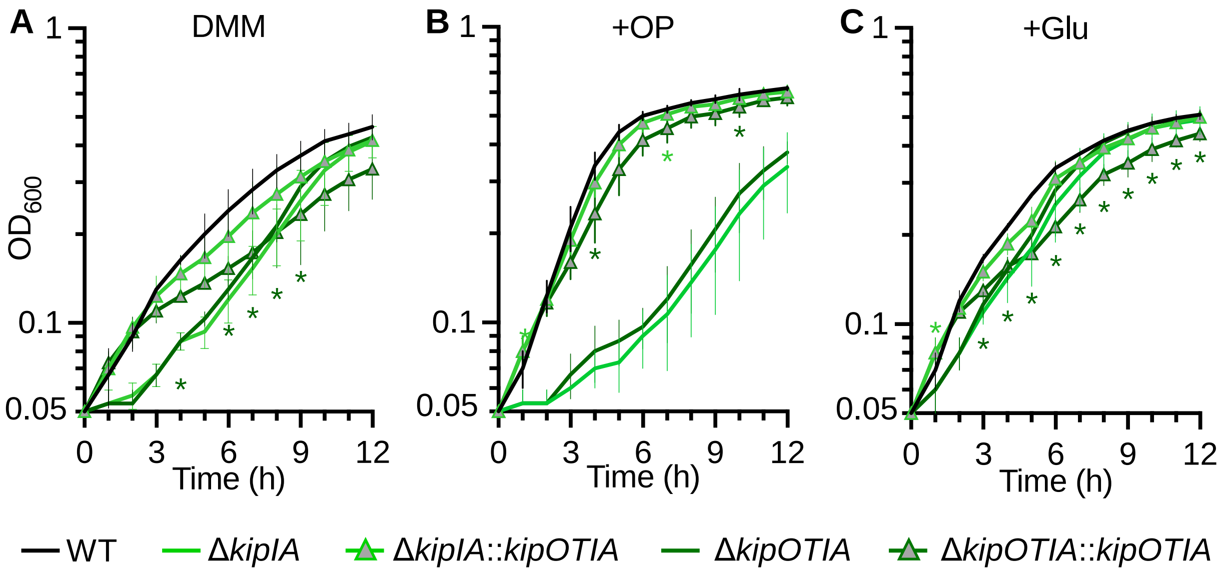


**Figure S5. Complementation improves the growth of *kip* mutants.** Condition-centered display of growth for 630Δ*erm* (WT), Δ*kipIA* (MC1903), Δ*kipOTIA* (MC2375), and complements Δ*kipIA*::*kipOTIA* (MC2519) and Δ*kipOTIA::kipOTIA* (MC2520) in **A)** DMM, **B)** DMM with 30 mM 5-oxoproline (+OP), and **C)** DMM with 30 mM glutamic acid (+Glu). The means and SD of three independent experiments are shown. Data were analyzed by one-way ANOVA with Dunnett's multiple comparison test at each timepoint for the respective complements to the wild-type. *, $P$ <0.05.

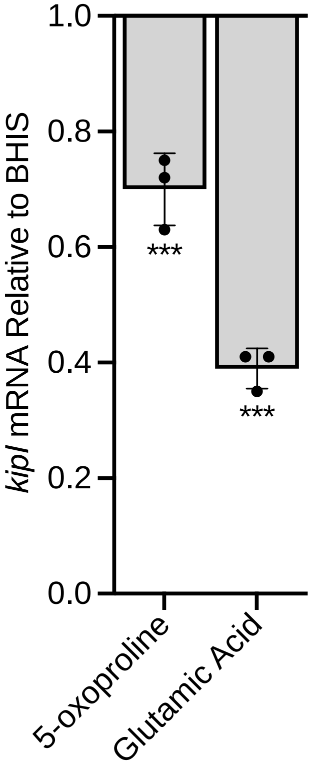


**Figure S6. Glutamic acid and 5-oxoproline modestly repress *C. difficile kipI* expression.** WT (630Δ*erm*) was grown in BHIS +/- 30 mM neutralized 5-oxoproline or 30 mM glutamic acid. Graph shows the mean, individual data points, and standard deviations of *kipI* mRNA expression levels relative to WT in BHIS alone for three independent experiments. Data were analyzed by one-way ANOVA with Dunnett's multiple comparison test comparing the treated WT with untreated WT. *** $P$ <0.001.
