## Supplemental Tables for "KipOTIA detoxifies 5-oxoproline and promotes the growth of *Clostridioides difficile*"

**Table S1.** Bacterial Strains and plasmids

| Plasmid or Strain | Relevant genotype or features | Source, construction, or reference |
|---|---|---|
| **Strains** | | |
| ***E. coli*** | | |
| DH5α Max Efficiency | F− Φ80*lac*ZΔM15 Δ(*lac*ZYA-*arg*F) U169 *rec*A1 *end*A1 *hsd*R17 (rk−, mk+) *pho*A *sup*E44 λ−thi−1 *gyr*A96 *rel*A1 | Invitrogen |
| HB101 | $F^-$ *mcrB mrr hsdS20*($r_B^-$ $m_B^-$) *recA13 leuB6 ara-14 proA2 lacY1 galK2 xyl-5 mtl-1 rpsL20* (conjugation) | B. Dupuy |
| ***C. difficile*** | | |
| 630Δ*erm* | $Erm^S$ derivative of strain 630 | N. Minton (Hussain et al., 2005, p. 630) |
| MC310 | 630Δ*erm spo0A*::*erm* | (Edwards et al., 2014) |
| MC324 | 630Δ*erm* pMC123 | (Edwards et al., 2014) |
| MC1903 | 630Δ*erm* Δ*kipIA*::*erm* | This study |
| MC1970 | MC1903 Δ*kipIA* pMC123 | This study |
| MC2375 | 630Δ*erm* Δ*kipOTIA*::*erm* | This study |
| MC2519 | 630Δ*erm* Δ*kipIA::kipOTIA*::*aad9* | This study |
| MC2520 | 630Δ*erm* Δ*kipOTIA::kipOTIA*::*aad9* | This study |
| ***B. subtilis*** | | |
| 1A1 | Wild-type, strain 168 lineage | (Koo et al., 2017) |
| BKE04080 | Δ*kipI*::*erm* | (Koo et al., 2017) |
| BKK04090 | Δ*kipA*::*erm* | (Koo et al., 2017) |
| MC2624 | BKE04080 → 1A1; Δ*kipI*::*erm* | This study |
| MC2625 | BKK04090 → 1A1; Δ*kipA*::*erm* | This study |
| **Plasmids** | | |
| pMSR | Pseudo-suicide plasmid used for allelic exchange in *C. difficile* strain 630; P*tet-CD2571.1 catP* | (Peltier et al., 2020) |
| pMC123 | *E. coli-C. difficile* shuttle vector; *bla, catP* | (McBride and Sonenshein, 2011) |
| pRT1099 | pMC123 with *aad9* cassette in place of *cat* | (Purcell et al., 2017) |
| pJIR1457 | shuttle vector with *ermBP* cassette | (Lyras and Rood, 1998) |
| pMC1076 | pMSR with *kipIA* homology regions flanking *ermB* | This study |
| pMC1233 | pMSR with *kipOTIA* homology regions flanking *ermB* | This study |
| pMC1354 | pMSR with homology flanking *kipOTIA* and *aad9* | This study |

**Table S2.** Oligonucleotides

| Primer | Sequence (5'→3')[a] | Use/locus tag/reference |
|---|---|---|
| oMC44 | CTAGCTGCTCCTATGTCTCACATC | *rpoC* (*CD0067*) qPCR; (McBride and Sonenshein, 2011) |
| oMC45 | CCAGTCTCTCCTGGATCAACTA | *rpoC* (*CD0067*) qPCR; (McBride and Sonenshein, 2011) |
| oMC501 | TGAGATTCCAGTATGTTATG | Forward primer in *kipI* (*CD1386*) cds |
| oMC502 | CTCTGCTTGTGTGTATTT | Reverse primer in *kipI* (*CD1386*) cds |
| oMC504 | TCTACAACCCATTCTATCT | Reverse primer in *kipA* (*CD1387*) cds |
| oMC2802 | GTAGAAATACGGTGTTTTTTGTTACCCTAAGTTTAAACGTGGAGCTGCACTTGGT | Forward primer for region 5' of *kipI* for Gibson assembly into pMSR |
| oMC2803 | TAATCTCATGACCAAAATCCCTTAACGAAATGTCACCTCTTCTAAATTAAGCAAATAAAG | Reverse primer for region 5' of *kipI* with homology to *ermB* |
| oMC2804 | CTTTATTTGCTTAATTTAGAAGAGGTGACATTTCGTTAAGGGATTTTGGTCATGAGATTA | Forward primer to amplify *ermB* cassette with homology to region 5' of *kipI* |
| oMC2805 | TTATAATAAAAATTGTATTTGTTAAAATTAAACTCCTTGGAAGCTGTCAGTAGTATACC | Reverse primer to amplify *ermB* cassette with homology to region 3' of *kipA* |
| oMC2806 | GGTATACTACTGACAGCTTCCAAGGAGTTTAATTTTAACAAATACAATTTTTATTATAA | Forward primer to amplify region 3' of *kipA* with homology to *ermB* |
| oMC2807 | GATTTTGGTCATGAGATTATCAAAAAGGAGTTTAAACTGGAGGTTCTTATGCTAGGTGAA | Reverse primer to amplify region 3' of *kipA* Gibson assembly into pMSR |
| oMC2820 | GTAGCTCTTGGAGGATTAGC | Forward primer to screen for *kip* deletions |
| oMC2823 | TCTACTATGTATTCTTATTGCCACTAC | Reverse primer to screen for *kip* deletions |
| oMC2868 | CAACATGTAAAACCACATG | Forward primer in *kipO* cds |
| oMC2869 | TAATCCACTTAATACAGTTCC | Reverse primer in *kipT* cds |
| oMC2870 | GCCACAATGATGAGAG | Forward primer in *kipT* cds |
| oMC2871 | ACGACGGCCAGTGAATTCATATGTACAAATATTGTGCATGTTTTAAAATAATATTTTAAC | Forward primer to amplify 500 bp 5' *kipO* and Gibson assemble into pMC123 |
| oMC3163 | AGGTCGACTCTAGAGGATCCTTAAACTGCTATATCATTTAATACTTTTGCCTTTA | Reverse primer in 3' end of *kipA* and Gibson assembly into pMC123 |
| oMC3343 | GTAGAAATACGGTGTTTTTTGTTACCCTAAGTTTAAACTACCAGCCAAAACAGAGCG | Forward primer to amplify 5' of *kipO* and Gibson assembly into pMSR |
| oMC3344 | TAATCTCATGACCAAAATCCCTTAACGTTTATGACCCCTCCTCGCTAA | Reverse primer to amplify 5' of *kipO* with homology to *ermB* |
| oMC3345 | TTAGCGAGGAGGGGTCATAAACGTTAAGGGATTTTGGTCATGAGATTA | Forward primer to amplify *ermB* cassette with homology to 5' *kipO* ( |
| oMC3538 | CTTTGTCTGGAGATAGTACAATAACA | Forward primer upstream of *kipO* to screen for *kip* deletions |
| oMC3585 | GTAGAAATACGGTGTTTTTTGTTACCCTAAGTTTAAACTTGTCTGGAGATAGTACAATAACACCT | Forward primer to amplify 5' of *kipO* region and Gibson assemble into pMSR |
| oMC3586 | CCAGTCACGTTACGTCGACTTAAACTGCTATATCATTTAATACTTTTGCCTTTAC | Reverse primer to amplify 5' of *kipO* with homology to *aad9* cassette |
| oMC3587 | GTAAAGGCAAAAGTATTAAATGATATAGCAGTTTAAGTCGACGTAACGTGACTGG | Forward primer to amplify *aad9* cassette with homology upstream of *kipO* |
| oMC3588 | CTATTTGTAAGTTATATTATAAATTATAATAAAAATTGTATTTGTTAAAATTAAAGTCGACACCCAAAATTGAAAAAAGTG | Reverse primer to amplify *aad9* cassette with homology 3' of *kipA* |
| oMC3589 | CACTTTTTTCAATTTTGGGTGTCGACTTTAATTTTAACAAATACAATTTTTATTATAATTTATAATATAACTTACAAATAG | Forward primer to amplify 3' of *kipA* with homology to *aad9* cassette |

| | | |
|---|---|---|
| oMC3590 | GATTTTGGTCATGAGATTATCAAAAAGGAGTTTAAACGCCACTACTTCTATGTTGTGCTATAATTAC | Reverse primer to amplify 3' of *kipA* to Gibson assemble into pMSR |
| oMC3966 | AGTCACGACGTTGTAAAACGACGGCCAGTGAATTCGTGGAGCTGATATAAGTGGAGC | Forward primer to confirm *B. subtilis* Δ*kipI* and Δ*kipA* mutants |
| oMC3973 | AGCTTGCATGCCTGCAGGTCGACTCTAGAGGATCCGCACGCACCAATTCAATTAAGAG | Reverse primer to confirm *B. subtilis* Δ*kipI* and Δ*kipA* mutants |

[a]Restriction sites underlined

**Table S3. Vector construction**

| Plasmid | Construction details |
|---|---|
| pMC1076 | A 909 bp homology arm 5' of *kipI* (*CD1386*) and a 796 bp homology arm 3' of *kipA* (*CD1387*) were amplified with primers oMC2802/2803 and oMC2806/2807, respectively. A 1523 bp *ermB* cassette from pJIR1457 was amplified with primers oMC2804/2805, and all three fragments were Gibson assembled into pMSR via the PmeI site. |
| pMC1233 | A 708 bp homology arm 5' of *kipO* (*CD1384*) and a 796 bp homology arm 3' of *kipA* (*CD1387*) were amplified with primers oMC3343/oMC3344 and oMC2806/2807, respectively. A 1523 bp *ermB* cassette from pJIR1457 was amplified with primers oMC3345/2805, and all three fragments were Gibson assembled into pMSR via the PmeI site. |
| pMC1354 | An 817 bp homology arm 5' of *kipO* (*CD1384*) and the 3772 bp *kipOTIA* operon (*CD1384 – 1387*) were amplified with primers oMC3585/3586 for a total fragment size of 4589 bp. An 864 bp homology arm 3' of *kipA* (*CD1387*) was amplified with primers oMC3589/3590. A 1045 bp *aad9* (spectinomycin resistance) cassette from pRT1099 was amplified with primers oMC3587/oMC3588. All three fragments were Gibson assembled into pMSR via the PmeI site. |
| pMC1181 | pMC123 with 500bp 5' of *kipOTIA* to encompass the native promoter for the *kipOTIA* operon and *kipOTIA.* pMC123 was digested with EcoRI and BamHI and the 500bp 5' of *kipOTIA* and *kipOTIA* fragment was amplified with oMC2871 and oMC3163. The two fragments were Gibson assembled into pMC123 via the EcoRI and BamHI sites. |

**Table S4. Composition of Defined Minimal Media (DMM)**

| Component | Final Concentration (mg $ml^{-1}$)[b] |
|---|---|
| **Amino Acids** | |
| L-Alanine | 0.535 |
| L-Arginine | 1.045 |
| L-Aspartic acid | 0.799 |
| L-Cysteine | 0.727 |
| L-Glycine | 0.450 |
| L-Histidine | 0.931 |
| L-Isoleucine | 0.787 |
| L-Leucine | 0.787 |
| L-Lysine | 0.877 |
| L-Methionine | 0.895 |
| L-Phenylalanine | 0.991 |
| L-Proline | 0.691 |
| L-Serine | 0.631 |
| L-Threonine | 0.715 |
| L-Tryptophan | 1.225 |
| L-Valine | 0.703 |
| L-Tyrosine[a] | 1.087 |
| **Salts** | |
| $Na_2HPO_4$ | 5.0 |
| $NaHCO_3$ | 5.0 |
| $KH_2PO_4$ | 0.9 |
| NaCl | 0.9 |
| **Trace salts** | |
| $(NH_4)_2SO_4$ | 0.00800 |
| $CaCl_2 \cdot 2H_2O$ | 0.0270 |
| $MgCl_2 \cdot 6H_2O$ | 0.0200 |
| $MnCl_2 \cdot 4H_2O$ | 0.0100 |
| $CoCl_2 \cdot 6H_2O$ | 0.00100 |
| **Iron** | |
| $FeSO_4 \cdot 7H_2O$ | 0.00400 |
| **Vitamins** | |
| D-Biotin | 0.00100 |
| Calcium-D-pantothenate | 0.00100 |
| Pyridoxine | 0.00100 |
| **$ZnCl_2$** | 0.0100 |
| **Sodium selenite** | 0.000179 |

[a]Tyrosine dissolved separately in $H_2O$ and HCl, then added after other amino acids
[b]Adjust media pH to 7.4 and filter sterilize

**Table S5. Composition of Modified Niehaus Defined Media**

| Component | Final Concentration (mg $ml^{-1}$)[a] |
|---|---|
| **Amino Acids** | |
| L-Tryptophan | 51.06 |
| **Salts** | |
| $K_2HPO_4 \cdot 3H_2O$ | 1.84 |
| $KH_2PO_4$ | 0.6 |
| $MgSO_4 \cdot 7H_2O$ | 0.5 |
| $MgCl_2$ | 0.0572 |
| **Trace salts** | |
| $CaCl_2 \cdot 2H_2O$ | 0.00735 |
| $MnCl_2 \cdot 4H_2O$ | 0.00099 |
| $CoCl_2 \cdot 6H_2O$ | 0.0006 |
| $ZnCl_2$ | 0.0017 |
| $CuCl_2 \cdot 2H_2O$ | 0.00043 |
| $Na_2MoO_4$ | 0.00052 |
| **Iron-Solution** | |
| $FeCl_3 \cdot 6H_2O$ | 0.02703 |
| $Na_3$-Citrate$\cdot 2H_2O$ | 0.088 |

[a]Adjust media pH to 7.0 and filter sterilize (Niehaus et al., 2017).
